# Diversification and conservation of DNA binding specificities of SPL family of transcription factors

**DOI:** 10.1101/2024.09.13.612952

**Authors:** Miaomiao Li, Tao Yao, Mary Galli, Luigi F. Di Costanzo, Wanru Lin, Yilin Zhou, Jin-Gui Chen, Andrea Gallavotti, Shao-shan Carol Huang

**Author notes:** Notice: This manuscript has been authored by UT-Battelle, LLC under Contract No. DE-AC05-00OR22725 with the U.S. Department of Energy. The United States Government retains and the publisher, by accepting the article for publication, acknowledges that the United States Government retains a non-exclusive, paid-up, irrevocable, worldwide license to publish or reproduce the published form of this manuscript, or allow others to do so, for United States Government purposes. The Department of Energy will provide public access to these results of federally sponsored research in accordance with the DOE Public Access Plan (http://energy.gov/downloads/doe-public-access-plan).

## Abstract

GRAPHICAL ABSTRACT

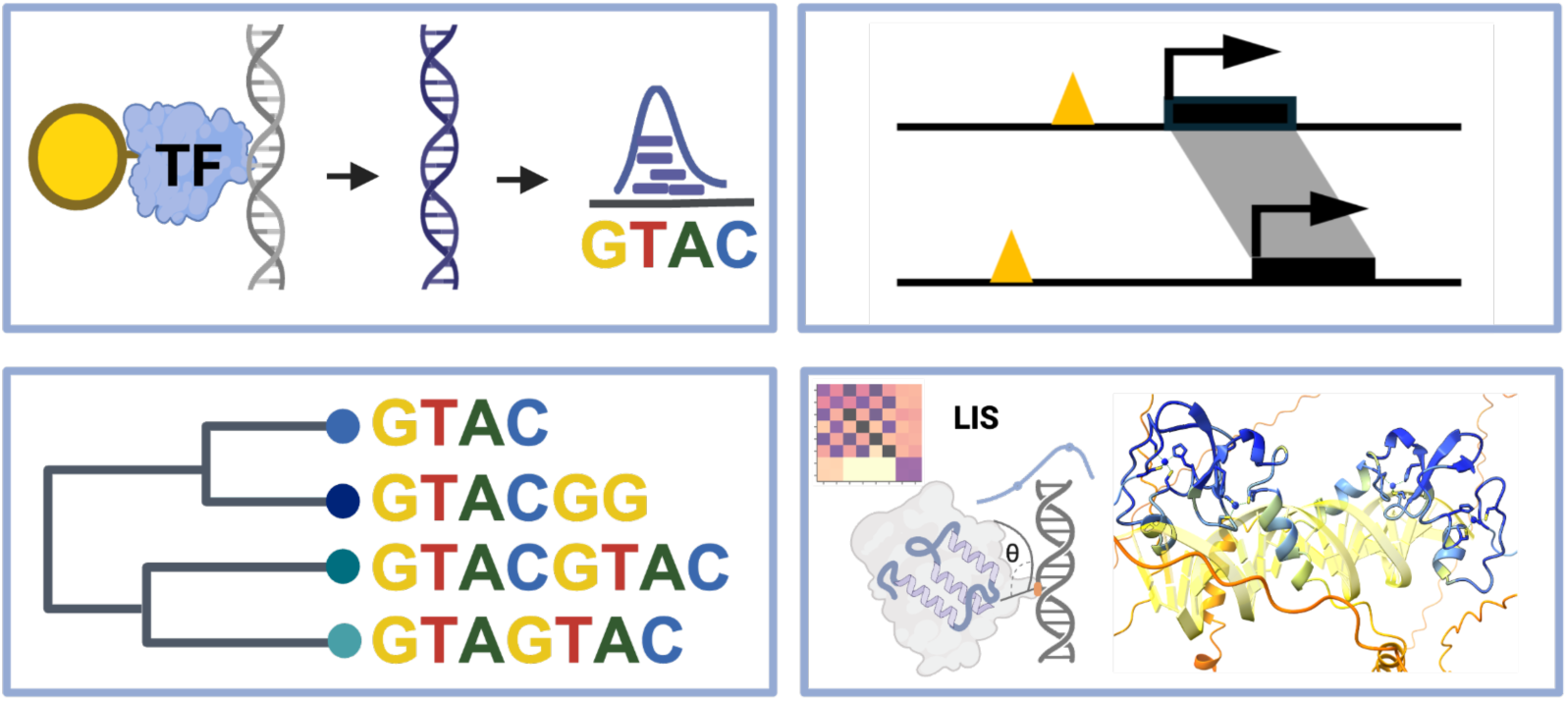

SQUAMOSA Promoter-Binding Protein-Like (SPL) transcription factors are key regulators of plant development and stress responses. Here, we present a comprehensive DNA affinity purification sequencing (DAP-seq) analysis of 14 of the 16 SPL members in *Arabidopsis thaliana*, revealing two major classes based on distinct DNA-binding motifs. Gene Ontology enrichment of target genes indicated both shared and specialized functions among AtSPLs, including roles in hormone signaling, metal ion homeostasis, and developmental transitions. Comparative analysis of closely related paralogs, AtSPL9 and AtSPL15, uncovered divergence in their genomic binding locations and target gene regulation, particularly in auxin and abscisic acid pathways. These differences were supported by motif analysis and protoplast reporter assays. Extending our study across species, we analyzed DAP-seq data from maize (*Zea mays*) and wheat (*Triticum aestivum)* SPL homologs, identifying conserved target genes but species-specific DNA motifs. Notably, wheat SPLs preferentially bind to longer, palindromic motifs distinct from the GTAC core motif in *Arabidopsis*. Using AlphaFold3 structure modeling, we show that these motif differences may arise from enhanced protein-protein interactions that stabilize dimeric binding. Our results highlight how SPL family members diversify through DNA-binding evolution and suggest that transcription factor dimerization contributes to motif complexity and regulatory specialization across plant genomes.

## INTRODUCTION

Transcription factors (TFs) are crucial regulators of gene expression, playing a central role in controlling various biological processes by binding to specific DNA sequences and modulating the transcription of target genes. The expansion of TF families through gene duplication events and subsequent functional diversification are a key evolutionary mechanism that enhances the complexity of gene regulation in plants (1). This expansion allows for the development of novel regulatory networks and the fine-tuning of gene expression in response to environmental and developmental cues. This process, known as functional diversification, can result in the emergence of TFs with altered DNA-binding specificities, thereby contributing to the evolution of complex regulatory circuits (1–3). A comprehensive analysis of DNA-binding specificities across an entire TF family is a powerful way to understand the full range of their regulatory potential. This can reveal how different TFs within a family contribute to the regulation of shared and distinct biological pathways, offering insights into the mechanisms underlying the evolution of gene regulatory networks.

As a plant-specific TF family, *SQUAMOSA PROMOTER BINDING PROTEIN-LIKEs* (SPLs) have highly conserved SQUAMOSA Promoter-Binding Protein (SBP) domains, which are approximately 76 amino acids long and contain two zinc finger-like structures and a nuclear localization signal motif (4). The first two SBP proteins discovered, *AmSBP1 a*nd *AmSBP2*, were identified in *Antirrhinum majus* floral meristem and found to act during early flower development (5). In *Arabidopsis*, the SPL genes are central to the age-dependent pathway of flowering (6–8). Many SPL members in *Arabidopsis* are targeted by miR156/157, and this miR156/SPL module plays an important role in diverse development stages including shoot meristem growth, flower development and the phase change from vegetative growth to reproductive growth, as well as the response to abiotic environmental stress (9–12). SPLs in *Arabidopsis* directly activate the expression of critical flowering genes, such as *SUPPRESSOR OF OVEREXPRESSION OF CONSTANS 1 (SOC1)*, *FLOWERING LOCUS T (FT)*, *FRUITFULL (FUL)*, and *APETALA1 (AP1)* (6, 13, 14), as well as auxin related genes. They also repress the expression of *AUXIN RESPONSE FACTOR (ARF)* genes by binding to their promoters such as those of *ARF6* and *ARF8*, thereby balancing auxin signaling (15).

In *Arabidopsis*, *AtSPL9* is one of the most extensively studied members of the SPL family (6, 7, 16). *rSPL9* is a miR156-insensitive mutant, showing reduced juvenile period and less response to aging (6, 17, 18). Within the *Arabidopsis* SPL family, *AtSPL9* and *AtSPL15* are the closest paralogs resulting from genome duplication and have redundant biological functions (19, 20). The *spl9/spl15* double mutants exhibit alternations in branching and flowering patterns, highlighting the critical role of AtSPL9 and AtSPL15 in regulating shoot branching (21). However, AtSPL15 and AtSPL9 coordinate distinct flowering phenotypes under short-day conditions (22). SPL homologs in crops play critical roles in controlling plant architecture, flowering time, and reproductive development. In rice (*Oryza sativa*), one of the most studied SPL genes, *OsSPL14*, also known as *Ideal Plant Architecture1 (IPA1*) and closely related to *AtSPL9* and *AtSPL15*, is a key regulator of tiller number and panicle size (8, 23). In maize (*Zea mays*), *ZmSBP8* and *ZmSBP30* are homologous to *AtSPL9/15*. The *zmsbp8/30* mutants exhibit notable changes in kernel row number, ear size, and cob architecture, ultimately affecting grain yield (24, 25). In hexaploid bread wheat (*Triticum aestivum*), the *AtSPL9/15* homologs across the three subgenomes are *TaSPL7A/B/D* and *TaSPL13A/B/D.* CRISPR/Cas9 knockout of *TaSPL7A/B/D* and *TaSPL15A/B/D* resulted in increased tiller number, decreased spike length, and reduced spikelet number (23, 26). These proteins are thus essential regulators of agronomic traits that directly impact crop yield, so identifying and editing the targets of the TFs could provide powerful tools for crop improvement.

Despite extensive research on the SPL family, key questions remain about the differences in DNA binding specificities among paralogs and the functional similarities and differences of these genes across plant species, especially in crops. In this study, we used DNA affinity purification-sequencing (DAP-seq) (27, 28), to map the genome-wide binding sites of the entire SPL family in *Arabidopsis* and systematically analyze their DNA binding specificities. Additionally, we utilized additional DAP-seq datasets for homologous SPLs in maize and wheat to examine the conservation of potential target genes SPLs across the three species, uncovering changes of their DNA binding specificities that could be mediated by a dimerization mechanism. By focusing on the expansion of the SBP/SPL TF family, our study provides insights into how TF duplication and subsequent functional diversification, mediated by evolution of DNA binding specificities, have contributed to the complexity of gene regulation in plants.

## MATERIALS AND METHODS

### Plant materials

*Arabidopsis* reference accession Col-0 (CS70000) was grown in soil at 21 °C under long-day (16 h light and 8 h dark) conditions for four weeks. Rosette leaves were collected and flash frozen in liquid nitrogen prior to genomic DNA (gDNA) isolation for DAP-seq DNA library preparation.

### DAP-seq experiments

The *Arabidopsis* SPL DAP and ampDAP-seq experiments were performed according to published protocols of (27, 29). First, frozen leaf tissues were ground into a fine powder using a mortar and pestle, dissolved in extraction buffer (0.1 M TRIS base, 0.5 M sodium chloride, 0.05 M EDTA, pH 8.0), and subject to phenol-chloroform extraction to remove proteins and other contaminants, resulting in purified gDNA. DAP-seq gDNA library was then prepared as a standard DNA sequencing library for the Illumina platform. gDNA was fragmented to an average of 200 bp using Covaris S220 Sonicator. Fragmented gDNA was end-repaired using the End-It DNA End-Repair Kit (Lucigen, ER81050, ordered in 2021) and incubated at room temperature for 45 min, followed by an A-tailing reaction using Klenow (3′→5′ exo-) (NEB, M0212) incubated at 37 °C for 30 min. A-tailed DNA fragments and annealed adapters were ligated in a ligation reaction using T4 DNA Ligase (Promega, M1804) incubated at room temperature for 3 h. The DNA was precipitated and suspended in elution buffer (10 mM Tris-Cl, pH 8.5), resulting in the gDNA libraries used in DAP-seq. A SPL protein expression reaction was prepared using TNT SP6 Coupled Reticulocyte Lysate System (Promega, L4600) incubated with 1000 ng pIX-HALO-SPL plasmids for 3 h at 30 °C. 10 μl of Magne HaloTag Beads (Promega, G7282) and 50 μl of wash buffer (PBS with 0.05% NP40) were added to the reaction, bringing the final volume to 100 μl, and the mixture was incubated on a rotator for 1 hour at room temperature. The proteins immobilized on beads were then washed five times on a magnet with 100 μl wash buffer to purify HaloTag-fused SPL protein. After washing, the protein-bound beads were incubated with 100 ng adapter-ligated gDNA library in 100 μl wash buffer for 2 h on a rotator. The mixture was then washed five times with wash buffer to remove unbound ligated DNA fragments. The beads were suspended in 30 μl elution buffer, heated at 98 °C for 10 min, and put on ice immediately for more than 5 min to denature the protein and release the bound DNA fragments. 25 μl of the supernatant was used for the PCR enrichment step. The ampDAP library is generated by amplifying DNA from the gDNA library by PCR, followed by incubation with the purified TF protein as for the DAP-seq protocol. The PCR reactions for making ampDAP-seq libraries and sequencing libraries following protein binding were prepared as 1 μl of Phusion DNA Polymerase (New England Biolabs, M0530), 10 μl of 5x Phusion HF Buffer, 2.5 μl of 10 mM dNTPs, 1 μl each 25 μM index primers, 25 μl of eluted DNA, add water to 50 μl. The PCR condition is as follows: 98 °C for 2 min, 15-19 cycles of 98 °C for 15 s, 60 °C for 30 s, and 72 °C for 1 min, final extension at 72 °C for 10 min. PCR-amplified samples were run in 1% agarose gel and cut to purify fragments from 200 bp to 600 bp with Zymoclean Gel DNA Recovery Kit (Zymo Research, D4007). The purified DNA fragments were measured by Qubit HS dsDNA Assay Kit (ThermoFisher, Q32854) and sequenced on the Illumina platform.

### Reporter assay in *Arabidopsis* protoplasts

To generate the effector plasmids used in the reporter assay, *AtSPL9* and *AtSPL15* CDS fragments were cloned in *pUC19-35S-DC* using LR reactions. A 758 bp sequence from the AtDI21 promoter (Supplementary Table S1) was synthesized by Twist BioScience and subcloned into *pUC19-DC-GUS* as reporter plasmid. The combinations of effector plasmid, reporter plasmid, and reference plasmid (*pUC19-35S-LUC*) were transformed into *Arabidopsis* leaf protoplasts. Middle sections of four-week-old fully expanded leaves were cut out, sliced into strips and immersed into enzyme solution containing 0.4 M mannitol, 20 mM KCl, 20 mM MES, 10 mM CaCl2, 5 mM β-mercaptoethanol, 0.1% BSA, 0.4% macerozyme R10, and 1.5% cellulase R10. The mixture was incubated at room temperature for 2 h before filtering the protoplasts through a 75 μm nylon mesh and washing them with W5 solution (154 mM NaCl, 125 mM CaCl2, 5 mM KCl, and 2 mM MES). After centrifuging at 1000 rpm for 3 min at 4 °C, the protoplasts were resuspended with MMg solution (0.8 M mannitol, 1 M MgCl2 and 0.2 M MES) to obtain a concentration of 200,000 cells per ml. Next, 6 μg effector, 3 μg reporter and 100 ng reference plasmids were co-transfected into 100 μl of protoplasts using the PEG-calcium mediated transfection method, followed by incubation in darkness for 18 to 20 h at room temperature. The GUS activity assay was conducted (30) and measured using a Fluoroskan microplate reader (30). The MUG (4-Methylumbelliferyl β-d-glucuronide) (Sigma-Aldrich, M9130) and luciferase assay system (Promega, E1500) were used to perform GUS and LUC activity assays, respectively. Relative GUS activity was calculated via normalization to LUC activity, and the data are presented as three independent biological replicates.

### DAP-seq data processing and analysis

#### Read processing, peak calling and motif discovery

The *Arabidopsis* DAP-seq libraries were sequenced on an Illumina NextSeq 500 instrument. The reads were basecalled using Picard IlluminaBasecallsToFastq (31) version 2.23.8, with APPLY_EAMSS_FILTER set to false. Following basecalling, the reads were demultiplexed using Pheniqs (32) version 2.1.0. The entire process was executed using a custom nextflow pipeline, GENEFLOW (33). Adapter sequences were trimmed from the single read FASTQ files by Trim Galore (version 0.6.6) and Cutadapt version 3.1 (34) with quality cutoff of 20. The trimmed reads were mapped to the *Arabidopsis* reference genome sequence TAIR10 using Bowtie2 (35)version 2.2.9 with default parameters. Aligned reads were filtered by mapping quality (MAPQ) score of at least 30 using SAMtools (36) version 1.11. Peak calling was done by the GEM peak caller (37) version 3.3 on the filtered mapped reads with the default read distribution, TAIR10 nuclear chromosome sequences, q-value threshold of 0.01 (--q 2), and parameters “--f SAM--t 1--min 200--k_min 5--k_max 14--k_seqs 2000--k_neg_dinu_shuffle -- outNP -- outBED -- outMEME -- outJASPAR -- outHOMER -- print_bound_seqs -- print_aligned_seqs”. Peaks were called for each replicate individually or by merging the two replicates using GEM’s multi-replicate mode, with samples from experiments of empty vector pIXHALO as control. To create a blacklist of regions that contained highly enriched but artifact signals, peak calling was done for a set of six pIXHALO empty vector controls using MACS3 (38), which could find broader regions of read enrichment compared to the point-source binding events reported by GEM. The peaks called by MACS3 (version 3.0.0a5) for these control samples with parameters “--keep-dup auto--nomodel--extsize 200 -q 0.1” were merged and the peak regions shared by at least two of these control samples were designated to be blacklist regions. DAP-seq peaks overlapping these regions were removed prior to all downstream analysis. BigWig files of normalized read signals were created using the MAPQ 30 filtered alignment BAM files by the bamCoverage program in the deepTools package (39) version 3.5.0 with the following parameters: “--binSize 1--normalizeUsing RPKM--ignoreForNormalization Mt Pt”. The read normalized bigwig files were visualized in JBrowse 2 (40). For motif discovery, the top 1000 binding events for each TF were obtained by sorting the GEM output narrowPeak files first by increasing q-value then by decreasing signal value. Sequences of 200 bp centered at the peak summits were extracted from the TAIR10 genome sequence and used as input for MEME motif discovery (41) with a Markov background model of order 2 computed from the peak sequences and parameters “-mod zoops-nmotifs 5-minw 5-maxw 15-dna-searchsize 0-revcomp-csites 1000”. The resulting PWM motifs were imported into R by the universalmotif package (42), aligned by the DiffLogo package (43), and plotted by the ggseqlogo package (44). The maize DAP-seq peak dataset, with blacklist regions already removed, was obtained from Zenodo under DOI 10.5281/zenodo.14991916 (45). The wheat DAP-seq data were obtained from NCBI GEO accession GSE188699 (26). Adapter sequences were trimmed from the pair-end FASTQ files by Trim Galore (version 0.6.6) and Cutadapt version 3.1 (34). The trimmed pair-end reads were mapped to the *T. aestivum cv. Chinese Spring v2.1* genome sequence assembly (46) using Bowtie2 (35) version 2.2.9 with default parameters. Aligned reads were filtered by MAPQ score of at least 20 using SAMtools (36) version 1.11. Peak calling was done by the MACS3 peak caller (38) version 3.0.0a5 with input library as the control and parameters “-f BAMPE -g 11453938932.0--keep-dup auto--nomodel--call-summits -q 0.05”. Blacklist regions of artifactual read enrichment were created by calling peaks on two replicates of the input libraries individually, merging the peaks and computing the regions called in both replicates of the input. DAP-seq peaks overlapping with blacklist regions were removed prior to all downstream analysis.

#### Arabidopsis DAP-seq sample clustering

MANorm2_utils (version 1.0.0) (47) were first used to find the number of DAP-seq reads of all *Arabidopsis* AtSPL DAP-seq samples contained in the GEM peak regions that occurred in at least one replicate. MAnorm2 R package (47) version 1.2.2 was used on this count matrix to perform normalization by a hierarchical normalization approach: the replicates for each TF were first normalized, then all the TFs were normalized to a pseudo-reference, a mean-variance curve was then fitted using parametric regression and occupied regions only. Peak regions ranked among the top 2,000 intervals by normalized signal in more than one sample were used to compute pairwise distances between TFs using the distBioCond function. The pairwise distance matrix was then used in hierarchical clustering of TFs with complete linkage method.

### Peak annotation and Gene Ontology Enrichment Analysis

To associate *Arabidopsis* DAP-seq peaks to genes, the top 3000 merged replicate GEM peaks for each TF were annotated with a gene that had the closest transcription start site (TSS) using the annotatePeak function in the ChIPseeker package (48) with the protein coding genes in Araport11 (49). Genes that had a DAP-seq peak within −1000 bp upstream and 500 bp downstream from the TSS were designated potential target genes. The R package clusterProfiler (50) was used to identify the top 10 most enriched GO terms for the target genes of each TF in the Biological Process ontology annotated in the org.At.tair.db database, and the enrichment P-values of these GO terms for all the TFs were computed. The enrichment P-values were corrected by the Benjamini & Hochberg method and a matrix of −log10 adjusted P-values was computed with the enriched GO terms on the rows and DAP-seq TFs on the columns and plotted as a heatmap by the ComplexHeatmap package (51). Clustering dendrogram was obtained by hierarchical clustering of the −log10 adjusted P-value matrix with Euclidean distance between rows and columns as distances and the complete linkage method.

### *spl7* RNA-seq data processing and Gene Set Enrichment Analysis

RNA-seq gene counts matrix for *spl7* mutant and Col-0 wild-type roots in low copper treatment and control conditions were downloaded from NCBI GEO accession GSE104916 (52). Using the R package DESeq2 (53), the count matrix was imported into R, pre-filtered by keeping only genes that had read counts more than 10 in at least 3 replicates, normalized and analyzed for differential expression with the design formula genotype+treatment+genotype:treatment. Differentially expressed genes between *spl7* and Col-0 wild-type (WT) in control condition were extracted from the contrast for the genotype factor between *spl7* and WT, with adjusted P-value threshold of 0.1. The genes were then ranked by the log2 fold change between WT and *spl7* and compared to the DAP-seq predicted target genes by the Gene Set Enrichment Analysis test (GSEA) function in clusterProfiler (50).

### Analysis of differential binding between *Arabidopsis* AtSPL9 and AtSPL15

Differential binding analysis between *Arabidopsis* AtSPL9 and AtSPL15 DAP-seq samples was performed by the R package DiffBind (54) version 3.6.5. The samples were normalized by the native DESeq2 normalization method RLE using reads in peaks, and differentially bound peaks were calculated by DESeq2 with the AtSPL9 as the reference and q-value threshold of 0.05. The significantly differentially bound peaks that have had a positive fold change were designated as AtSPL15-preferred and those that had a negative fold change were designated as AtSPL9-preferred. Sequences were extracted from the TAIR10 reference genome for the top 1000 AtSPL15-preferred or AtSPL9-preferred peaks sorted by adjusted P-values for motif discovery using MEME-ChIP (55) version 5.3.0 with the parameters “-meme-mod anr-meme-searchsize 0-minw 5”. Genes were associated with the top 3000 AtSPL15-preferred or AtSPL9-preferred peaks and GO enrichment was performed as described above.

### Cross-species comparison of AtSPL9/15 and homologs in maize and wheat

DAP-seq peaks for maize and wheat were annotated using the R package ChIPseeker (48), with the Zm00001eb.1 genome annotation for maize (56) and the IWGSC RefSeq v2.1 annotation for wheat (46). Genes with DAP-seq peaks located within 10,000 bp upstream and 500 bp downstream of the TSS were designated as potential target genes. Homologs between maize and *Arabidopsis* and between wheat and *Arabidopsis* were taken from the Best.hit.arabi.name column in the PhytozomeV13 annotation_info file for maize and wheat, respectively (57). Conserved target genes between maize and *Arabidopsis* were defined as genes associated with peaks shared between ZmSBP8 and ZmSBP30 that are homologous to genes associated with peaks shared between AtSPL9 and AtSPL15. For wheat and *Arabidopsis*, conserved targets were defined as genes associated with peaks shared by at least two of TaSPL7A/B/D and at least two of TaSPL13A/B/D that were homologous to those genes associated with peaks shared between AtSPL9 and AtSPL15. To identify DNA-binding motifs, sequences within the DAP-seq peaks associated with conserved target genes were extracted from their respective genomes. Motif discovery was then performed using MEME-ChIP with the parameters “-ccut 0-meme-mod anr-minw 4-meme-nmotifs 5”.

### Structure modeling of SPL-DNA interaction

To investigate the structural basis of differential DNA motif binding among SPL homologs from *Arabidopsis*, maize and wheat, we used AlphaFold3 (AF3) (58) to predict structures of the SPL TFs interacting with DNA sequences containing the different motif types. Our goal was to determine whether PPIs could explain why SPL homologs in different species prefer distinct DNA motifs, i.e., GTAC for *Arabidopsis* SPLs and GTASTAC for wheat SPLs, by evaluating differences in TF-TF and TF-DNA interaction potential. To this end, we modeled both one-copy and two-copy TF-DNA complexes for each protein in combination with either its species-preferred binding motif or the motif preferred by an orthologous SPL in a contrasting species.

For protein inputs, we used full-length amino acid sequences corresponding to the isoforms analyzed in the DAP-seq experiments (26, 45). For DNA inputs, we focused on DAP-seq peaks associated with conserved SPL target genes across species. Within these peaks, we scanned for motif matches corresponding to the preferred motif of the SPL in question, for example, GTAC in *Arabidopsis* SPL peaks and GTASTAC in wheat SPL peaks. We selected the ten highest-scoring motif matches for each protein as representative binding sites. For each selected site, we extracted a 20 bp DNA sequence centered on the motif match, yielding a set of DNA fragments we refer to as “species-preferred motif sequences”. To create a comparison set, we then searched genomic regions nearby (within 10 kb of the preferred motif) for high-scoring matches to the cross-species motif, i.e., the motif preferred by a homologous SPL in another species. Importantly, these cross-species motif matches were selected to not overlap any DAP-seq peaks, to reduce the likelihood that they are bound by the SPL under study. We extracted 20 bp DNA fragments centered on these motifs to create the “cross-species motif sequences”. Thus, for each SPL protein, we generated two matched DNA input sets: one representing motifs preferred and bound by this SPL and one representing motifs not preferred or bound by this SPL but preferred by a homologous SPL in another species.

Each protein-DNA combination was then modeled using AF3 with at least six replicate prediction runs using different random seeds. Each run used either a one-copy or two-copy version of the TF sequence paired with one of the twenty 20 bp-long DNA sequences including the forward and reverse complement (RC) strands, as well as two zinc ions for each copy of the protein. The resulting structures were visualized using ChimeraX v1.10.1 (59). Confidence in structural predictions was assessed using AF3’s standard metrics: predicted template modelling (pTM) score, the interface predicted template modeling (ipTM) score, and predicted aligned error (PAE). Local Interaction Score (LIS) calculations were performed by modifying the Jupyter notebook provided using the recommended parameters (60). For each combination of protein (one-or two-copy) and motif (species-preferred or cross-species motif) condition, we computed the median LIS score across all replicate runs for each of the ten DNA sequence instances. Wilcoxon rank-sum tests were used to compare the LIS score distributions of the ten motif matches in the species-preferred motif sequences versus the cross-species motif sequences.

## RESULTS

### Genome-wide binding site analysis of *Arabidopsis* SPL family by DAP-seq

We applied DAP-seq to all 16 members of the SPL family in *Arabidopsis,* successfully identifying between 854 to 34,301 peaks (read enrichment q-value ≤ 0.01) for 14 members (Figure 1A). By using ampDAP-seq, which assays protein binding on a PCR-amplified DNA library, we obtained 2,189 peaks for AtSPL16 (Figure 1A). With the exception of AtSPL11 and AtSPL13, all other AtSPLs had thousands of peaks, indicating strong DNA binding activities. In contrast, AtSPL11 (854 peaks) or AtSPL13 (no peaks) may have potential DNA binding activities that are less detectable under the tested experimental conditions in DAP-seq, or they might require cofactors to enhance their DNA binding activity. The genome-wide binding site maps of the AtSPL members showed both conserved and variable binding signals at the promoter regions of the well-known downstream targets, for example, the flowering development gene *SOC1* and an auxin response gene *ARF8* (Figure 1B and 1C). The distribution of binding events relative to genomic features was similar for all the AtSPL members, with around 30-40% of the binding events located in proximal promoter regions of - 1000 bp to 500 bp from the transcription start sites (TSS) (Figure 1D).

**Figure 1:**
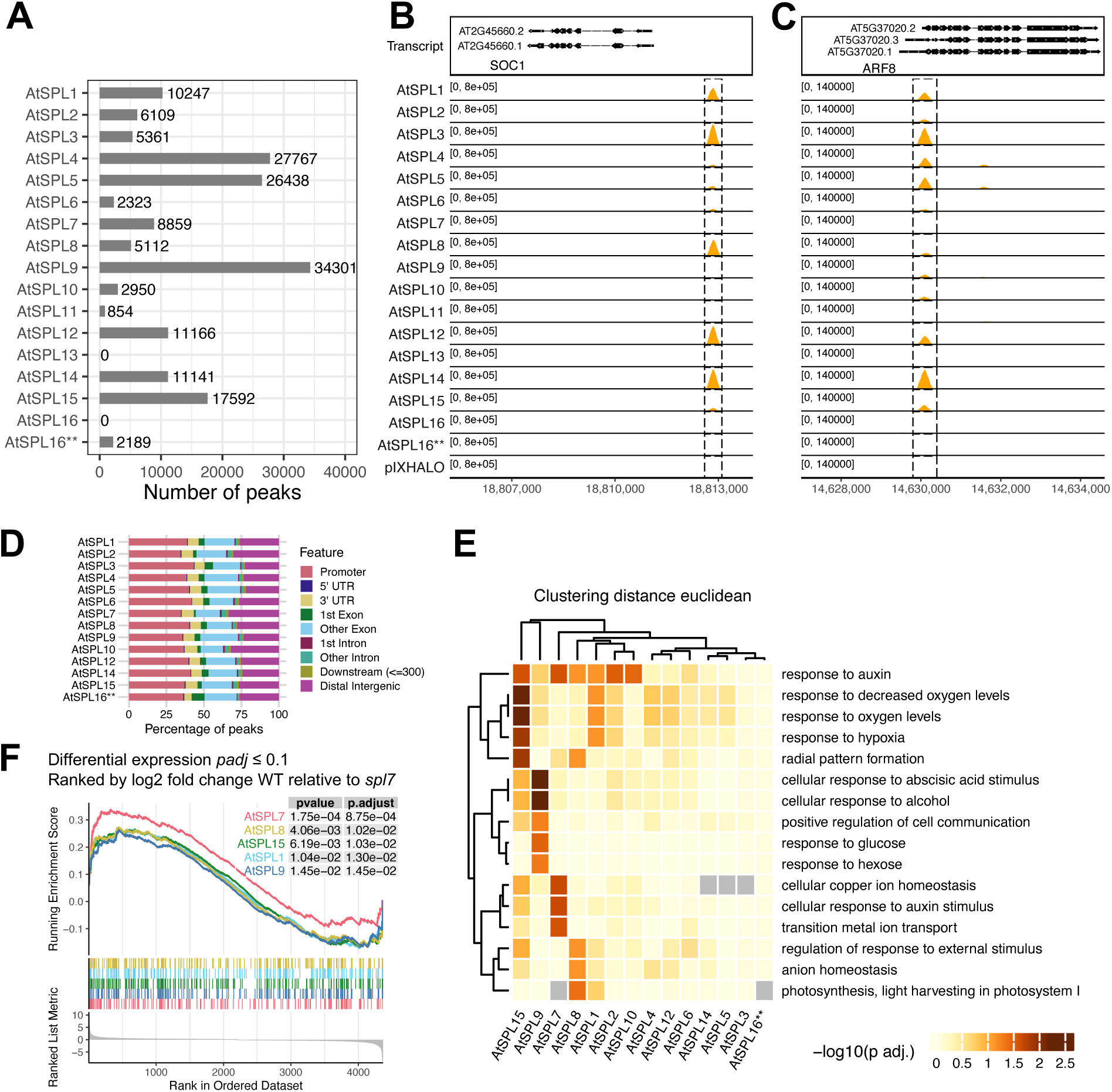
DAP-seq identifies genome-wide binding events for SPL TFs that are biologically relevant. (**A**) Number of DAP-seq peaks for all SPL members in *Arabidopsis*. ** indicates ampDAP-seq. (**B**) *Arabidopsis* AtSPL DAP-seq binding signal at the promoter region of a known target gene *SOC1*. (**C**) *Arabidopsis* SPL DAP-seq binding signal at the promoter of *ARF8*. (**D**) Distribution of AtSPL DAP-seq peaks at genome annotation features. Promoter is defined as - 1000 bp upstream to 500 bp downstream of TSS. (**E**) Enriched Gene Ontology biological process terms for DAP-seq predicted targets of *Arabidopsis* SPLs. (**F**) GSEA of AtSPL DAP-seq targets and the differentially expressed genes in *spl7* mutant vs. Col-0 wild type (WT).

To gain insight into the potential biological functions of the AtSPL family members based on their DNA binding profiles, we performed Gene Ontology (GO) enrichment analysis of genes associated with the top 3000 binding events ranked by DAP-seq binding site scores. The top ten enriched biological processes across family members revealed their shared and distinct functions (Figure 1E). Notably, hormonal responses, such as response to auxin and abscisic acid (ABA), were highly enriched for AtSPL15 and AtSPL9. Hypoxia and oxygen-related responses were more enriched for AtSPL15, AtSPL1, AtSPL2, AtSPL4 and AtSPL12. In contrast, the GO term “photosynthesis, light harvesting in photosystem I” was uniquely enriched for AtSPL8, suggesting a potential role in capturing light energy. Interestingly, metal ion homeostasis processes, such as cellular copper ion homeostasis and transition metal ion transport, were predominantly enriched for AtSPL7 DAP-seq targets, consistent with its previously characterized function in regulating copper uptake and distribution, which is essential for plants’ adaptation to low copper conditions (13, 52, 61, 62).

We next evaluated the potential for DAP-seq reported binding events to regulate gene expression by testing the association between the DAP-seq predicted target genes and differentially expressed genes (DEG) in *spl* mutants, in particular, in the root of *spl7* mutant from a published dataset (52). Using Gene Set Enrichment Analysis (GSEA) tests (63, 64), we found that among the AtSPL members that had strong GO term enrichment, DAP-seq targets of AtSPL7 were the most enriched for down-regulated DEGs between *spl7* mutant and wild-type Col-0 (up-regulated comparing wild-type and mutant; Figure 1F), suggesting that DAP-seq can predict transcriptionally regulated genes by the relevant TFs for further functional studies.

### Genome-wide binding and DNA sequence motifs support two major classes in the AtSPL family

To investigate the relationships between the *Arabidopsis* SPL family members in terms of their DNA binding specificities, we first clustered the AtSPLs based on their genome-wide binding patterns reported by DAP-seq. To do this, we first determined the regions of the highest DAP-seq read signals among all the AtSPL binding profiles and used the read enrichment signal in these regions to calculate the pairwise distances between the individual TFs (47). Hierarchical clustering of the distance matrix revealed two major subgroups (Figure 2A): Class A containing four SPLs (AtSPL2, 10, 7 and 11) and Class B containing the rest of the SPLs (AtSPL9, 15, 1, 12, 4, 14, 5, 6, 3, 8). We then performed *de novo* motif discovery for each AtSPL using sequences in the 1000 peaks with the highest significance of DAP-seq read enrichment in each experiment. We found that the most significant motifs recognized by the Class A members shared a core GTAC motif, flanked by a weaker T base on either side, while Class B members preferred a core GTACGG motif (Figure 2B). These results suggest Class A and Class B members could bind to distinct sets of genes. For example, the well-known gene *HSL2* (also known as *VAL2*), which promotes vegetative phase change (65), showed strong binding signals at its promoter region from all Class B members that preferred the GTACGG motif. In contrast, relatively low binding signals were observed for Class A members that preferred the GTAC motif (Figure 2C). Additionally, in Class B, motifs for AtSPL9 and AtSPL15 (close relatives derived from a duplication event) contained weaker GG consensus sequence adjacent to the GTAC core, suggesting these two paralogs bind DNA with more flexibility outside of the core motif than other Class B members.

**Figure 2:**
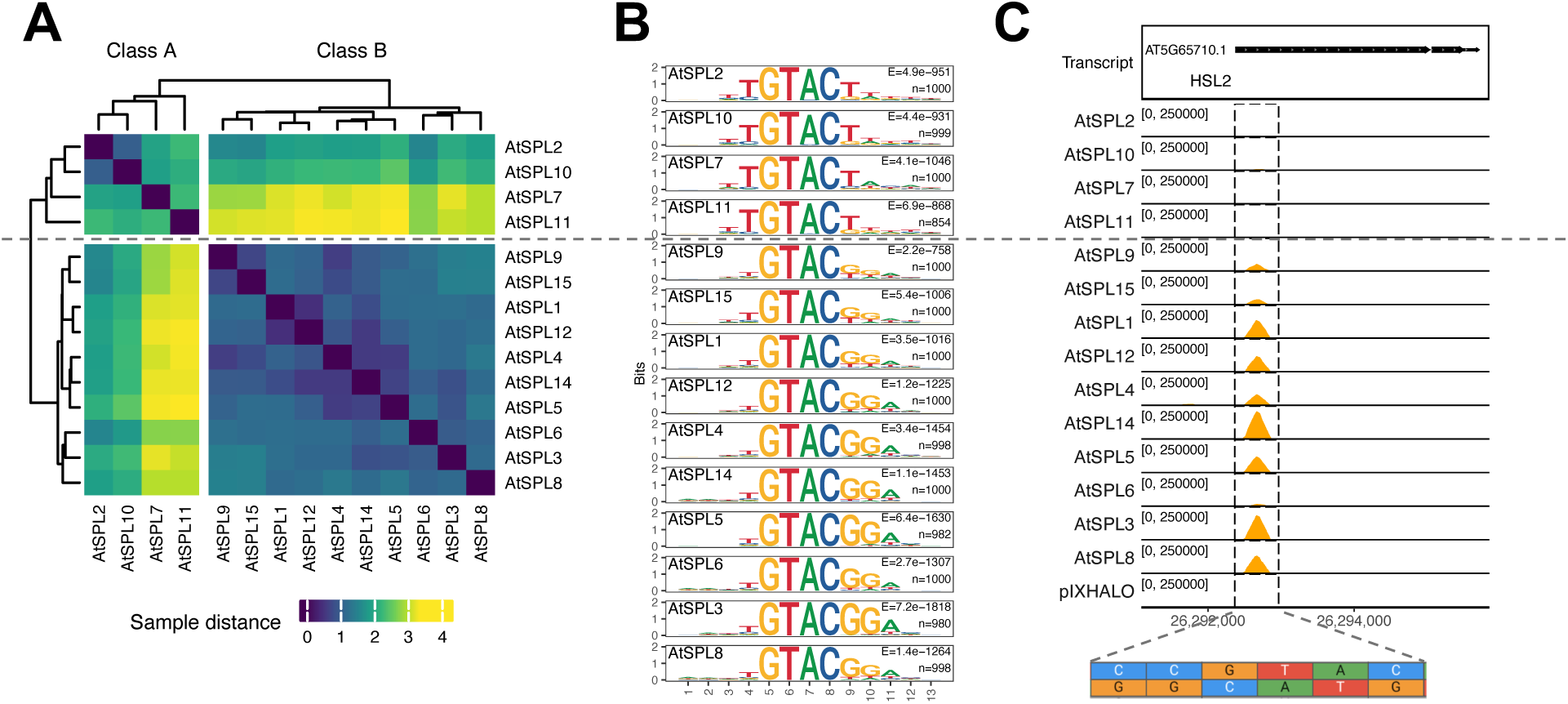
Genome-wide binding profile comparison and DNA motifs of the *Arabidopsis* SPL family. (**A**) Hierarchical clustering of AtSPL DAP-seq binding profiles splits the family members into Class A and Class B. (**B**) PWM models of the enriched motifs from the top 1000 DAP-seq peaks for each AtSPL. (**C**) Class A and Class B AtSPLs show different DAP-seq binding signals at the promoter of phase change related gene *AtHSL2* (*AtVAL2*).

### DAP-seq revealed differential DNA binding specificities between paralogs AtSPL9 and AtSPL15

To investigate the functional coherence or divergence of the evolutionarily conserved AtSPL9 and AtSPL15, we conducted differential binding analysis (54) between the AtSPL9 and AtSPL15 DAP-seq data (54). We identified 5,066 AtSPL15-prefered and 3,122 AtSPL9-prefered binding sites at FDR threshold of 0.05 (Figure 3A; Supplementary Table S2), about 18% and 15% of the binding sites for these two SPLs, indicating potential functional distinctions between them (Figure 3A). Strikingly, distinct biological functions were enriched for genes associated with the AtSPL15- and AtSPL9-preferred binding events (Figure 3B). AtSPL15-preferred gene targets were strongly enriched in processes related to ion homeostasis, positive regulation of transcription, response to oxygen and hypoxia, and auxin response. In contrast, AtSPL9-preferred gene targets were mainly enriched in secondary cell wall biogenesis, responses to photooxidative stress and regulation of the abscisic acid (ABA)-activated signaling pathway. These results suggest that beyond the conserved functions, AtSPL15 and AtSPL9 could have distinct roles in regulating different biological processes, and DNA binding preferences in the genome could mediate their specialized functions in plant development and stress responses.

**Figure 3:**
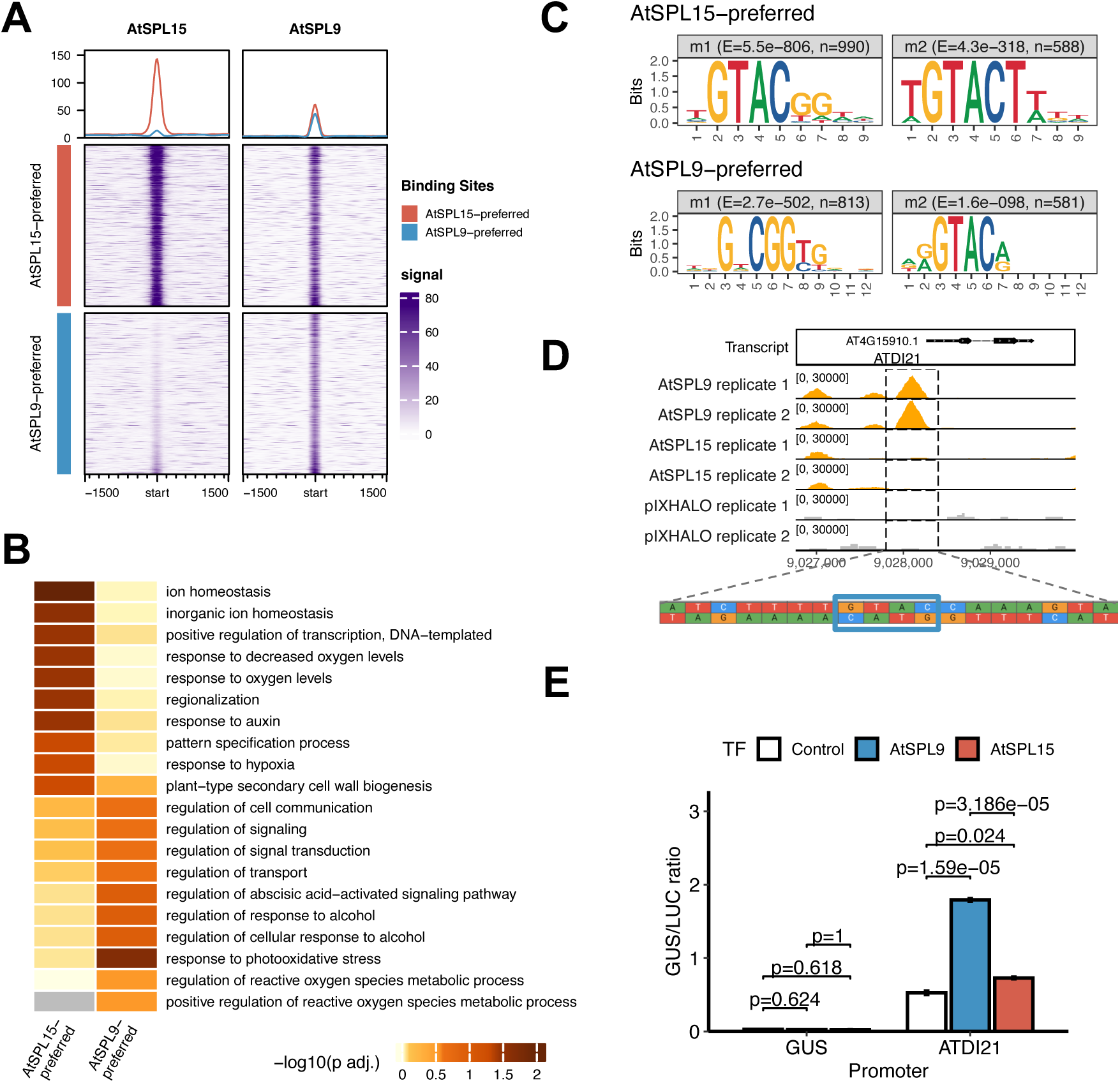
Differential binding events, target genes and motifs comparing *Arabidopsis* AtSPL9 versus AtSPL15. (**A**) DAP-seq read signals at differentially bound peaks of AtSPL15 and AtSPL9. (**B**) Top 10 enriched GO Biological Process terms for genes associated with AtSPL15-preferred and AtSPL9-preferred peaks. (**C**) PWM motif models enriched in AtSPL15- and AtSPL9-preferred binding sites. (**D**) AtSPL9-preferred binding site upstream of gene *AtDI21*. (E) Transient expression of AtSPL9 and AtSPL15 induced significantly different expressions of the reporter AtDI21::GUS. Error bars are standard error of the mean (n=3) and P-values were computed by two sample t-tests followed by Bonferroni correction.

To understand the mechanism underlying the differential DNA binding preferences between AtSPL9 and AtSPL15, we performed *de novo* motif discovery using the sequences in the 1000 differentially bound peaks with the highest increase in binding by AtSPL15 and AtSPL9, respectively (Figure 3C). We found that the most significant motifs in the AtSPL15-preferred binding sites contained the consensus sequences GTACgg and TGTACT, while AtSPL9 preferred motifs contained the consensus sequences GNCGG and GTAC. As an example, the differentially bound peak at the promoter region of the gene *DROUGHT-INDUCED 21* (*AtDI21*; AT4G15910) showed strong binding by AtSPL9 but not by AtSPL15, and it contained a binding site sequence GTAC (Figure 3D). To validate that the differential binding events could influence gene expression, we performed reporter assays (66) by cloning a 758 bp sequence from the promoter region of *DI21*, including the 200 bp AtSPL9-preferred DAP-seq peak, into the reporter plasmid *pUC-GUS* (*DI21pro::GUS*) to drive the GUS reporter gene expression. When we transiently expressed AtSPL9 and AtSPL15 by transfecting SPL9 and SPL15 separately, we found that AtSPL9 induced significantly higher reporter activation compared to AtSPL15 (Figure 3E). These results support our hypothesis that AtSPL15 and AtSPL9 have distinct binding preferences to DNA, resulting in differences in target gene regulation.

### Comparative analysis of SPL target genes in *Arabidopsis*, maize, and wheat suggests conserved functions

To characterize the DNA binding specificities among SPL homologs across plant species, we compared the conserved genes associated with DAP-seq peaks - referred to as “target genes” - of AtSPL9/15 in *Arabidopsis*, ZmSBP8/30 in maize (67) and TaSPL7/13 in wheat (including three paralogs from the A/B/D subgenome) (26) from DAP-seq experiments conducted on their respective genomes (26, 67). By comparing the target genes of maize ZmSBP8/30 to those of *Arabidopsis* AtSPL9/15, we identified 4,557 unique target genes for AtSPL9/15, 1,125 unique target genes for ZmSBP8/30, and 492 conserved target genes shared between the two species (Figure 4A; Supplementary Table S3). Similarly, when comparing wheat TaSPL7/13 target genes with the target genes of AtSPL9/15 in *Arabidopsis*, we found 4,017 unique target genes for *Arabidopsis* AtSPL9/15, 2,709 unique target genes for wheat TaSPL7/13, and 939 conserved target genes (Figure 4B; Supplementary Table S4).

**Figure 4:**
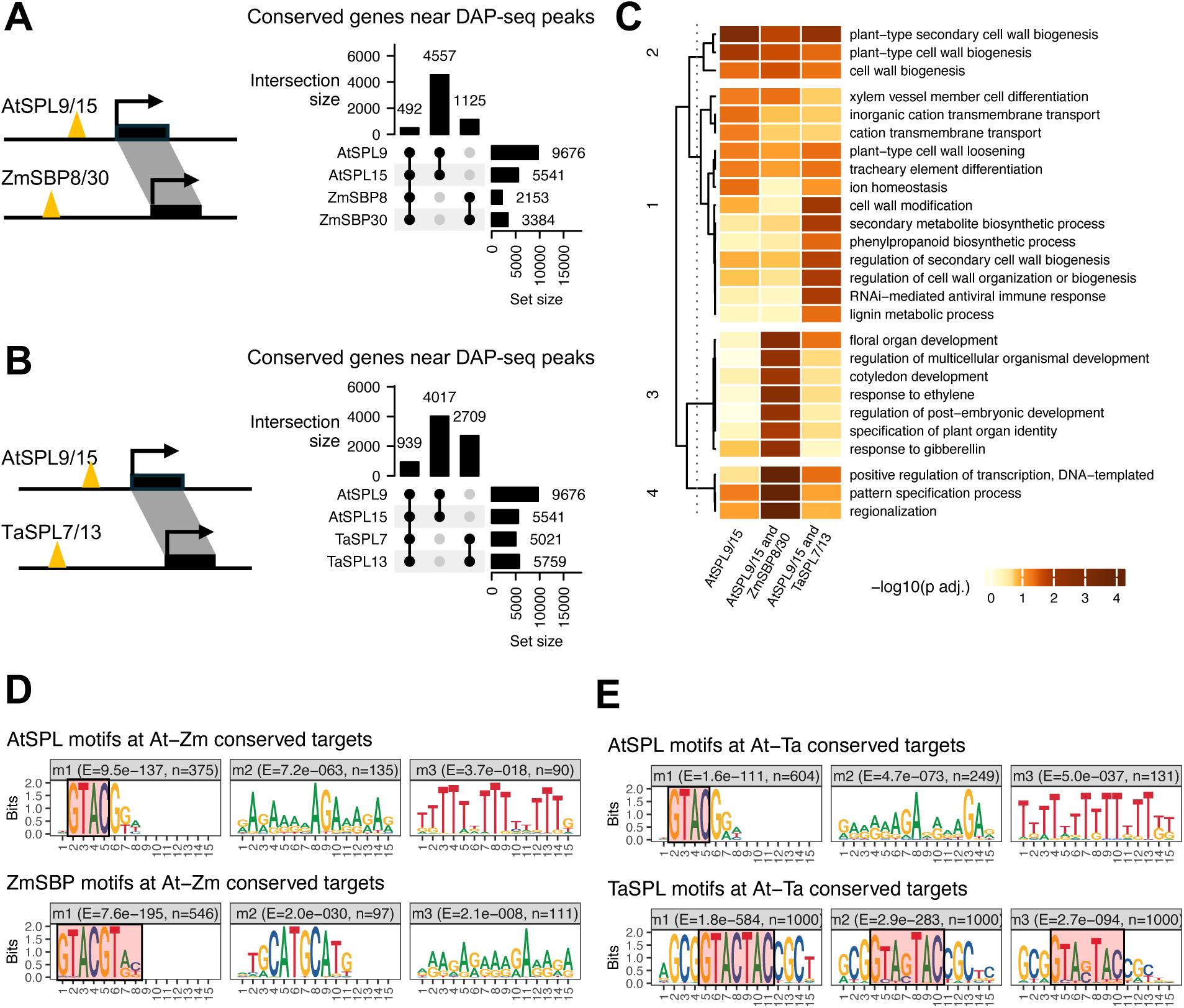
Comparative analysis of DAP-seq target genes and binding site sequences revealed variation of DNA binding mechanism among SPL homologs. (**A**) Upset plot comparing the target gene overlap between *Arabidopsis* AtSPL9/15 and maize ZmSBP8/30. (**B**) UpSet plot comparing the target gene overlap between *Arabidopsis* AtSPL9/15 and wheat TaSPL7/13. (**C**) GO enrichment analysis showing the biological processes enriched for targets conserved between SPL homologs between *Arabidopsis* and maize and between *Arabidopsis* and wheat. (**D**) DNA binding motifs discovered by MEME for DAP-seq peaks associated with conserved SPL targets in *Arabidopsis*, maize and wheat.

To gain insight into the function of these conserved target genes, we performed GO enrichment analysis and found four major clusters of enriched biological processes (Figure 4C). Cluster 2 and Cluster 4 highlighted functions conserved across the three species, primarily related to cell wall biogenesis, transcriptional regulation, and pattern specification. Cluster 1 also revealed functions that were conserved among all three species, such as xylem and cell wall development, while also suggesting *Arabidopsis*-wheat-specific conserved processes such as cell wall modification, secondary metabolite biosynthesis, and lignin metabolic processes. These GO terms suggest the potential role of TaSPL7/13 in specific aspects of wheat architecture, possibly related to structural integrity and stress responses. Cluster 3 GO terms were significantly enriched for SPL targets conserved between *Arabidopsis* and maize, including processes associated with different developmental stages, such as floral organ development, cotyledon development, and post-embryonic development. Additionally, we observed processes such as plant organ specification and gibberellin response, which are closely related to flowering time and overall plant development. Overall, these results suggest that different biological functions and developmental processes are targeted by the SPL shared target genes in different species, offering insights into the evolutionary trajectories of the SPL homologs in the three species spanning monocot to dicot.

### SPL homologs bind to variations of the GTAC motifs

Since conservation of binding motifs is often used as the basis for cross-species prediction of *cis*-regulatory elements, we analyzed the enriched DNA sequence motifs in the DAP-seq peaks associated with the conserved target genes in the three species. Unexpectedly, we identified three different types of DNA binding motifs, although they all shared or partially shared the core GTAC sequence (Figure 4D-E, red highlights). In the *Arabidopsis* genome, AtSPL9/15 recognizes motifs with a GTAC core adjacent to a GG dinucleotide. In the maize genome, ZmSBP8/30 recognizes a back-to-back repeat of the GTAC sequence, GTACGTRC, although one of the GTAC elements is weak. In the wheat genome, TaSPL7/13 binds to a motif containing two strong GTAC elements overlapping by one base pair, resulting in the consensus GTASTAC (GTACTAC or GTAGTAC). These different yet related motif architectures suggest differences in genome-wide DNA recognition mechanisms among the SPL homologs. Importantly, the more complex back-to-back or overlapping motifs in maize and wheat may contribute to enhancing binding specificities in the much larger and complex genomes of maize (2.3 Gbp) and wheat (17 Gbp) relative to *Arabidopsis* (135 Mbp).

### Structure modeling links protein-protein interaction preferences to DNA binding motif variation across SPL homologs

To explore the potential structural mechanisms underlying the motif variation among SPL homologs, we performed structural prediction of the SPL-DNA binding complexes using AlphaFold3 (AF3) (58). We began by modeling the interaction between full-length SPL proteins and DNA sequences containing their species-preferred binding motifs. These sequences were drawn from DAP-seq peaks associated with conserved target genes, representing binding events that are conserved across species and therefore likely to be functionally important. As shown in Figure 5A for the representative homologs, TaSPL13A interacting with the TaSPL-preferred GTASTAC motif and AtSPL9 interacting with the AtSPL-preferred GTAC motif, the predicted structures had low overall confidence scores (ipTM: 0.51-0.59; pTM: 0.29-0.31), likely due to high fractions of disordered regions outside the conserved SBP DNA-binding domain (fraction disordered: 0.80-0.81). Nevertheless, when we superimposed the AF3-predicted structures with the experimentally solved X-ray crystal structure of AtSPL5 (PDB 8J49 Chain A (68)), we observed strong structural similarity in the SBP domains, including two well-conserved zinc-binding pockets (RMSD 0.685-0.672 Å, 63 Cα atoms; Figure 5B). Furthermore, the predicted structures consistently positioned the zinc finger regions in the C-terminal portion of the SBP domain in proximity to the DNA (Figure 5B), consistent with prior solution studies showing that this zinc-binding site is required for DNA binding (69). While these results confirmed structural conservation of the SBP domain across species, they did not readily explain the observed motif divergence across species.

**Figure 5:**
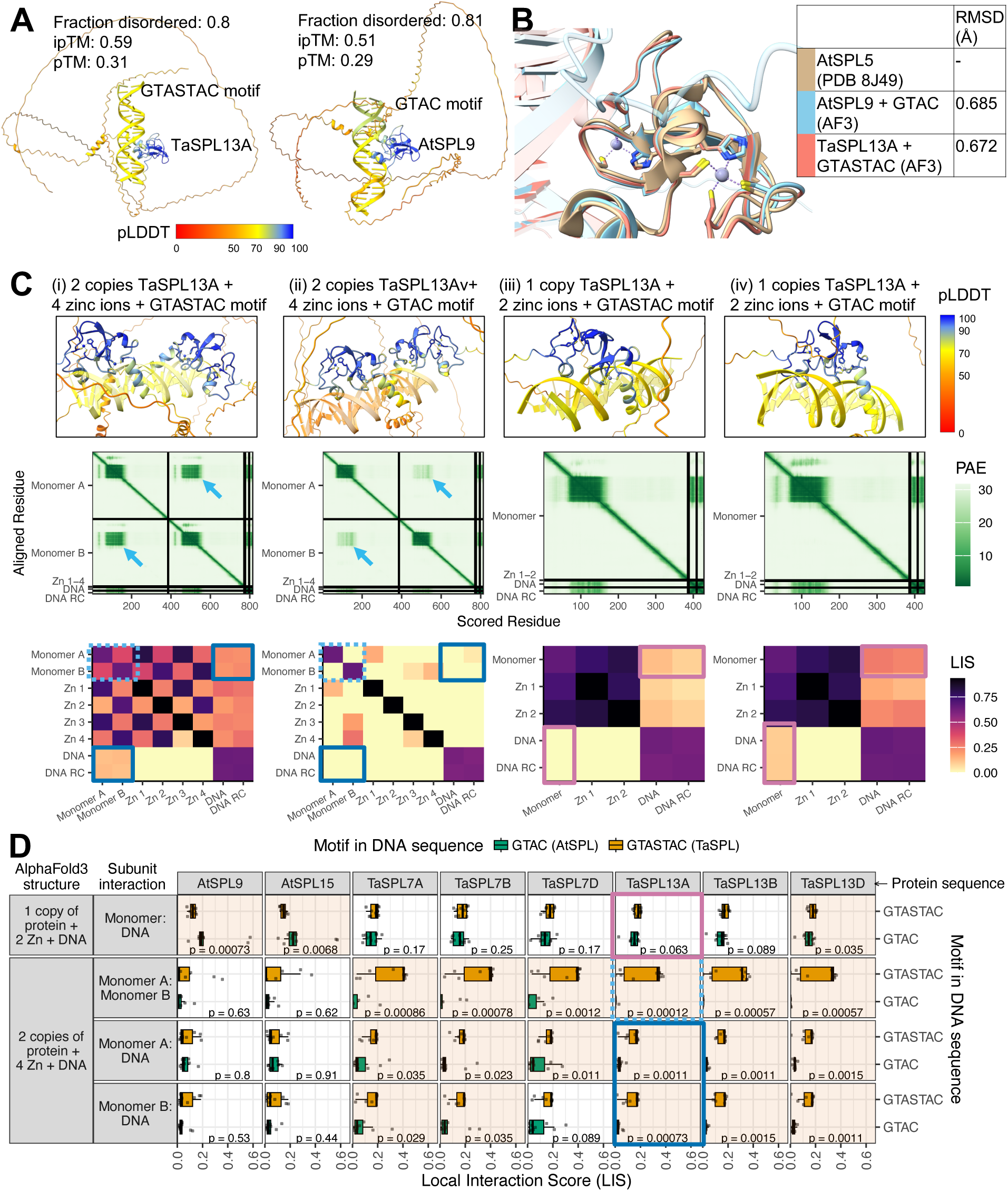
AlphaFold3 (AF3) structure modeling of DNA binding by *Arabidopsis* and wheat SPL proteins. (**A**) Representative AF3-predicted structures of TaSPL13A binding to its preferred DNA motif GTASTAC (left) and AtSPL9 binding to its preferred motif GTAC (right). (**B**) Superposition of the SBP domains from the predicted structures in (A) with the X-ray crystal structure of AtSPL5 (PDB: 8J49), highlighting the conserved zinc-binding regions. (**C**) AF3-predicted complex structures, Predicted aligned error (PAE) matrices, and Local Interaction Score (LIS) heatmaps for one or two copies of TaSPL13A interacting with either the species-preferred GTASTAC motif or the cross-species GTAC motif (see MATERIALS AND METHODS). Colored boxes highlight predicted monomer-monomer (cyan), monomer-DNA (blue) interactions in two-copy protein structures, and monomer-DNA interactions (purple) in one-copy protein structures. RC: Reverse Complement. (**D**) Summary of LIS values across predicted SPL-DNA structures. Each point represents the median LIS from n≥6 replicate predictions for a given motif instance. For *Arabidopsis* SPLs (AtSPL9 and AtSPL15), structures were modeled with one or two protein copies interacting with either GTAC motifs (species-preferred motifs bound in DAP-seq) or GTASTAC motifs (cross-species motifs, unbound). For wheat SPLs (TaSPL7A/B/D and TaSPL13A/B/D), structures were modeled analogously with GTASTAC (species-preferred motifs bound in DAP-seq) and GTAC (cross-species motifs, unbound) motifs. Boxplots summarize monomer-monomer and monomer-DNA interaction scores for each configuration. P-values were computed by two sample Wilcoxon rank-sum tests between species-preferred and cross-species motif conditions and orange-shaded panels indicate statistically significant differences (p < 0.05).

We next hypothesized that the wheat SPLs’ strong preference for the overlapping GTAC motifs (GTASTAC) may arise from enhanced protein-protein interactions (PPIs) between SPL monomers. This hypothesis was motivated by three observations. First, DAP-seq has been shown to capture dimeric protein binding to DNA for several TF families in multiple species (45, 70, 71). Secondly, DNA-guided TF interaction is known to produce overlapping or composite motifs that partially reflect specificities of individual TFs (70, 72). Third, dimerization has been observed in several SPLs in different species. In *Arabidopsis*, structural and biochemical studies show that the SBP domains of *Arabidopsis* SPLs bind DNA as monomers (69, 73, 74), while full-length proteins like AtSPL7 and AtSPL9 are capable of dimerization, though the functional role of this dimerization in DNA binding remains unclear (75, 76). In particular, AtSPL7 dimerization may modulate its regulatory activity *in vivo*, possibly by preventing its translocation from the cytoplasm to the nucleus (75). In maize, the SBP-box TF ZmTGA1 forms both monomeric and dimeric complexes to bind DNA. Notably, the dimeric form of ZmTGA1 exhibits higher stability than that of its teosinte ortholog, and this increased dimer stability was implicated as a key molecular mechanism underlying the naked kernel phenotype that distinguishes domesticated maize from its wild progenitor, teosinte (77). Together, these observations suggest that PPI-driven DNA binding could underlie the emergence of complex binding motifs in SPLs.

To test this hypothesis, we used AF3 to predict SPL-DNA complex structures with two copies of each SPL interacting with their preferred binding motif (species-preferred motif) or with a motif preferred by SPL homologs from a different species (cross-species motif) (see MATERIALS AND METHODS). These DNA sequences were drawn from genomic regions associated with conserved SPL target genes and centered on high-scoring motif matches either within a SPL’s own DAP-seq peaks (species-preferred motif sequences) or from nearby unbound regions containing the motif of the SPL homolog in a different species (cross-species motif sequences). This design allowed us to compare predicted structural interactions in sequence contexts that are biologically relevant while minimizing confounding effects from local genomic features. Figure 5C shows representative results for TaSPL13A, comparing its interaction with the species-preferred GTASTAC motif (structure i) versus the cross-species GTAC motif preferred by AtSPLs (structure ii). In the predicted aligned error (PAE) matrices, a strong low-error block was present between the two TaSPL13A monomers in the GTASTAC-bound complex but absent in the GTAC-bound complex (cyan arrows), suggesting a more stable dimer interface between the two TaSPLA monomers in the presence of the species-preferred motif.

Since AF3 is better at predicting the overall geometry of TF-TF-DNA complexes than localizing the precise TF-TF or TF-DNA contacts (70, 78), we did not analyze the direct contacts in the predicted structures. Instead, we calculated the local interaction score (LIS) between the subunits in the predicted structures (60). LIS has improved sensitivity in detecting PPIs, particularly for those that involve flexible and small interfaces where traditional pTM scores are less informative (60). The LIS heatmaps shown in Figure 5C confirmed that both monomer-monomer (cyan boxes) and monomer-DNA (blue boxes) interactions were stronger in the TaSPL13A complex bound to the GTASTAC motif (structure i) compared to that bound to the GTAC motif (structure ii). In contrast, control structures predicted using a single copy of TaSPL13A (structures iii and iv; Figure 5C) did not show differences in LIS values for the monomer-DNA interaction (purple boxes) that are consistent with the motif preferences, supporting the idea that PPIs between TaSPL13A monomers stabilize the binding to the GTASTAC motif.

To generalize these results, we similarly modeled one- and two-copy SPL-DNA complexes for AtSPL9, AtSPL15, six wheat TaSPL7/13 paralogs, and two maize ZmSBPs ZmSBP8 and ZmSBP30. For each protein, we predicted structures with both the species-preferred motif sequences and cross-species motif sequences (see MATERIALS AND METHODS), and computed the LIS values for the monomer-monomer and monomer-DNA interactions. The resulting LIS distributions across ten motif instances for species-preferred and cross-species motifs are shown as dotplots and boxplots in Figure 5D. Statistical comparisons between predictions for species-preferred versus cross-species motifs (Wilcoxon rank-sum test, p < 0.05; orange background panels in Figure 5D) revealed a consistent and striking pattern. *Arabidopsis* AtSPL9 and AtSPL15 showed significantly higher LIS values for the GTAC motif only in the one-copy models. In contrast, most wheat SPL paralogs exhibited significantly stronger monomer-monomer and monomer-DNA interaction scores when modeled with the GTASTAC motif, but only in the two-copy structures. These results support a model in which PPIs between TaSPL monomers enhance binding to the overlapping GTASTAC motif, whereas AtSPLs bind DNA as monomers with a preference for the short GTAC motif.

Interestingly, applying the same analysis to maize ZmSBP8 and ZmSBP30 did not show consistent preferences between one-copy and two-copy structures for either the GTAC or GTACGTRC motifs (AtSPL-preferred or ZmSBP-preferred) (Figure S1). This may reflect the flexibility of maize SBPs in forming monomeric or dimeric complexes, such as those found for ZmTGA1 (77).

## DISCUSSION

TFs regulate gene expression by binding to specific DNA sequences, and variations in their binding sites can lead to differential regulation of genes within a TF family in a species and across different species. Our analysis of the DAP-seq binding profiles of the *Arabidopsis* SPL family revealed two major classes of binding site motifs (Figure 2), yet the enriched GO terms of the target genes near the most enriched binding sites were much more diverse among the family members (Figure 1E). This suggests that even closely related TFs, such as AtSPL9 and AtSPL15, can develop distinct regulatory roles through subtle differences in their DNA-binding preferences. By analyzing this comprehensive cistrome dataset for SPL TF family in *Arabidopsis*, our study offers insights into how specific DNA-binding characteristics contribute to the functional diversity within the SPL family of TFs.

Gene duplication is a key feature of genomic architecture, especially within TF families, leading to opportunities for subfunctionalization and neofunctionalization (79–82). Some duplicated TFs retain ancestral functions by maintaining their original DNA binding preferences, while others acquire mutations that allow them to bind different DNA sequences and thus regulate different sets of genes (2, 83, 84). Such changes can enhance a species’ ability to respond to broader environmental challenges including biotic or abiotic stresses, or activate new developmental pathways for adaptation. DAP-seq offers a powerful approach to investigate this functional divergence mediated by DNA binding, allowing for detailed scrutiny of DNA binding motifs and target genes among paralogous TFs, even when genetic redundancy makes functional studies challenging. For example, our analysis revealed that even slight changes in DNA binding motifs between the duplicated homologs AtSPL9 and AtSPL15 led to distinct target gene sets, with AtSPL15 showing preferences for genes involved in auxin responses, while AtSPL9 favoring targets related to ABA signaling. These variations help differentiate the roles of closely related TFs in regulating downstream targets and ensuring functional specificity. We expect that the same analysis can be applied to other TF homologs within the SBP/SPL family or extend to other TF families to gain a broader understanding of TF functional diversity.

The evolution of TF-DNA interactions is a complex and dynamic process that involves multiple factors, including changes in the TF proteins, their DNA-binding motifs, and the genomic context of the binding sites such as DNA shape and chemical modifications (3). In this study, we observed that the conserved functions of the SPL/SBP homologs based on their DNA binding targets were distributed differently in *Arabidopsis*, maize, and wheat: while some functions maintained across all three species, others were shared only between specific pairs, such as *Arabidopsis* and maize or *Arabidopsis* and wheat. For example, common regulatory functions related to developmental processes were found between *Arabidopsis* and maize, while *Arabidopsis* and wheat shared functions in stress response and structural regulation. Analysis of DAP-seq binding sequences revealed species-specific preferences for related but distinct motif variants, which may contribute to the divergence of regulatory networks across species. These findings provide insights into how the SPL/SBP TFs have evolved to balance the core, conserved functions with species-specific adaptations, highlighting the power of DAP-seq to shed light on the evolutionary trajectories of TF networks across plant species.

A particularly interesting finding emerged when we compared the DNA binding motifs among the SPL/SBP homologs across species, revealing that the SPL/SBP homologs in maize and wheat bind to longer motif types that are variations of the core GTAC sequences recognized by the *Arabidopsis* protein. Specifically, the GTACGTRC motif in maize is a back-to-back palindromic sequence of two GTAC units, while the GTASTAC motif in wheat is a palindromic sequence with one base pair overlap between two GTAC units (Figure 4E). Palindromic motifs have also been observed for dimers of the bZIP family of TFs, suggesting such motifs can be a general mechanism for refining TF-DNA recognition (64). Many TF families, including AP2/ERF, bZIP, ARF, homeodomain and MADS-box, are known to form dimers or higher-order complexes to bind DNA and regulate gene expression (28, 85, 86). Using AF3 structural modeling, we found that TaSPL7/13 in wheat may recognize the palindromic GTASTAC motif through DNA-guided dimerization, enabling binding to a more complex motif configuration than the GTAC motif preferred by AtSPLs. The ability to recognize longer, more complex motifs through PPI could be a mechanism that allows SBP/SPL TFs to increase DNA binding specificities to achieve precise gene regulation, particularly in the context of complex plant genomes like those of maize and wheat. Our findings underscore the potential role of PPI in the evolution of DNA recognition mechanisms by TFs, offering insights into how TFs adapt to regulate diverse regulatory programs in plants with highly divergent genome complexities.

## AUTHOR CONTRIBUTIONS

S.C.H. and M.L.: Conceptualization, Formal analysis, Visualization, Methodology, Validation, Writing-original draft. W.L. and Y.Z.: Validation. M.G. and A.G.: Formal analysis. T.Y. and J.C.: Methodology, Validation. L.F.D.C.: Investigation, Methodology. S.C.H. and A.G.: Funding acquisition. All authors: review & editing.

## ACKNOWLEDGEMENTS

We thank Joseph Ecker for the gift of TF expression plasmids. We also thank Aurelia Li for technical assistance. We acknowledge the Zegar Family Foundation for their generous support. This work was supported in part through the NYU IT High Performance Computing resources, services, and staff expertise.

